# Electrospun Poly (ε-caprolactone) Membranes Modified with Heparin and Essential Fatty Acids for Biomedical Applications

**DOI:** 10.1101/2024.02.08.579539

**Authors:** Taisa Farias, Joelma Ricardo, Jessica Cunha, Yonny Barcelay, Ariamna Dip, Camila Ruzo, Ivanildes Bastos, Karen Segala, Joel Silva Junior, Ştefan Ţălu, Marco Paula, Walter Brito

## Abstract

The study aimed to enhance wound healing by modifying poly(ε-caprolactone) (PCL) membranes with sodium heparin (HS) and Essential Fatty Acids (EFA). Electrospinning was used to prepare the membranes containing the maximum concentration of HS and AGE, which were then sterilized with ozone. Microbiological tests confirmed effective sterilization. The membrane characterization included scanning electron microscopy (SEM) for morphological analysis, wettability studies for contact angle determination, Fourier-transform infrared spectroscopy (FT-IR) analysis, and thermogravimetry (TGA) for thermal analysis. Results indicated successful preparation and sterilization of PCL membranes modified with HS and EFA. Morphological analysis showed well-formed and randomly distributed fibers, although the PCL+HS membrane exhibited beads on its fibers. Adding HS and EFA affected the fiber diameter, with PCL+HS fibers having a smaller diameter than pure PCL and PCL+EFA fibers. Wettability analysis demonstrated modified surface properties with reduced contact angles. FT-IR analysis showed slight contributions of HS and EFA in the modified PCL membranes, while thermal analysis revealed no substantial changes in thermal stability. In conclusion, PCL membranes modified with HS and EFA can potentially accelerate wound healing, presenting an innovative and cost-effective approach to treating skin injuries.

## 1. Introduction

Developing products from biocompatible and bioabsorbable materials requires broad scientific knowledge in engineering, chemistry, biology, and biophysics. Engineering is crucial in transforming raw materials to create valuable products for humans, including developing polymeric materials applicable in the pharmaceutical and medical industries.^1^

Polymeric materials are highly versatile and widely used in various applications, such as water purification membranes, food packaging films, and conductive sensor elements. One significant application is the production of controlled drug release systems for biomedical applications. These systems aim to control the concentration of drugs temporally and spatially, maximizing clinical benefits and minimizing adverse effects.^2,3^

Electrospinning has stood out among the various techniques used to develop polymeric membranes. This technique allows the fabrication of continuous nanoscale fibers and has been extensively explored in biomedical areas, such as the production of three-dimensional porous scaffolds, controlled drug release systems, high-efficiency filters, protective clothing, catalysts, adsorbent materials, and sensors.^4^ Electrospun membranes of poly(ε-caprolactone) (PCL) are widely researched devices in tissue engineering technologies, as well as in studies of controlled drug release and coverage of complex wounds, such as burns and wounds in diabetic patients.^5,6^ A drug of interest for release in complex wounds is heparin, a sulfated polysaccharide with anticoagulant properties. Heparin is widely used as a therapeutic drug, and its association with biocompatible materials allows for controlled drug release^7^. Additionally, there are essential oils from Amazonian plants and other regions, such as sunflower oil, which possess wound healing and anti-inflammatory properties due to the presence of essential fatty acids (EFA) such as omega-9, omega-6, and omega-3.^8^

Reducing the healing time of wounds is of great economic and social importance, especially for public hospitals, as it reduces hospitalization time and associated costs. In this context, the production of electrospun PCL membranes containing natural and synthetic drugs shows promise due to their biocompatible, biodegradable, and non-toxic characteristics.^9,10^

## 2. Materials and Methods

### 2.1. Materials

PCL with a molar mass of 50,000 g/mol and 30,000 g/mol, sodium heparin at a concentration of 5,000 IU/mL, and sunflower oil rich in essential fatty acids EFA from the commercial brand Curatec AGE Essencial, which contains a high concentration of essential fatty acids, medium-chain triglycerides, and vitamins A and E, were used in the study. The solvents used were chloroform (99% purity) and acetone (99.8% purity), acquired from Biotec. The syringes used were of model SR 10 mL, and their needles had a diameter of 0.8 mm.

### 2.1. Methods

#### 2.1.1. Electrospinning

The process parameters were determined, such as the needle diameter (0.8 mm), volumetric flow rate (8 mL/h), and working distance (distance between the needle and the metallic collector of 18 cm). These parameters remained fixed for all electrospinning trials.^11^ It is essential to emphasize that the applied voltage to the polymeric solution is also a process parameter. Still, it cannot be kept constant as it depends on the surface tension of each solution. The environmental parameters, such as air humidity (%) and temperature, are recorded during the process.

For the PCL solution, the solvents were weighed on an analytical balance (METTLER TOLEDO model AB204) in a 1:1 mass ratio of solvents (equivalent to 3.16 g of each solvent). They were then dissolved using a magnetic stirrer (NOVA ÉTICA model 113) for 15 minutes. Subsequently, 1 g (13.66%) of PCL pellets were added and mechanically stirred for 16 hours in a closed system.

To study the maximum concentration of EFA in the PCL polymer solution, 4 electrospinning trials were conducted with 0.25 g, 0.50 g, 0.75 g, and 1 g of sunflower oil. The solutions were mechanically stirred for 4 hours.

Four experiments were performed using HS in the PCL polymer solution. The quantities of HS used were 0.10 mL, 0.20 mL, 0.30 mL, and 0.40 mL. The solutions were mechanically stirred for 8 hours.

#### 2.1.2. Sterilization using ozone

The sterilization process used a PODOXI ozone generator with a flow rate of 6.5 L/min and a maximum dosage of 150 ppm. The membranes were placed inside a desiccator with pressure control. Subsequently, a vacuum was applied using a pump, and for the aeration period, ozone gas was injected at the maximum concentration of the equipment, maintaining a continuous vacuum. This procedure was repeated four times with intervals of 5 minutes. After sterilization, the membranes were removed from the desiccator and subjected to vacuum packaging using a Registron vacuum sealer, model RG 300 A.

Three different culture media were employed for the sterility test: Agar-Agar, Sabouraud Agar, and Brain Heart Infusion (BHI) broth. Each culture medium was used to test for microorganisms, enterobacteria, pneumococci, streptococci, meningococci, fungi, yeasts, and aerobic and anaerobic bacteria. The process involved weighing the culture medium, hydrating it with distilled water, and sterilizing it with moist heat in an autoclave (121°C for 15 minutes). Afterward, the sterile culture medium was also autoclaved in previously sterilized Petri dishes.^12^

The test was performed in triplicate for each membrane type, using four Petri dishes in each case: one control dish and three others containing three sample discs for the three-culture media. The dishes were incubated at 37°C for 14 days in a bacteriological incubator with the lid facing down to avoid contamination of the culture media by condensation water.

#### 2.1.3. Morphological characterization

This stage was carried out using the TESCAN VEGA3 equipment. The membranes were gold-metalized using a BAL TEC CPD 050 metallizer to obtain the micrographs. It was possible to obtain measurements of fiber diameters from the acquired images. The measurements were performed manually using the ImageJ image processing software.

#### 2.1.4. Study of membrane wettability through contact angle

The wetting assay was performed using a digital microscope (DINO Lite plus) with a magnification capacity of up to 1000 times. At room temperature, a 10 μL deionized water droplet was deposited onto the membrane surface. The behavior of this droplet was observed for 120 seconds. Contact angle measurements were obtained with the assistance of Image J software.

#### 2.1.5. Chemical characterization by Fourier-transform infrared spectroscopy

The spectra in the infrared region with Fourier-transform (FTIR) were obtained using the Agilent Cary 630 FTIR equipment, with 128 scans in the range of 4000 to 650 cm^-1^ and a resolution of 8 cm^-1^.

#### 2.1.6. Thermal analysis of the membranes through Thermogravimetry

For the study of the thermal stability of the electrospun membranes obtained in this work, a thermogravimetric analysis - TGA was performed by heating up to 1000°C at a rate of 50°C/min under a nitrogen gas atmosphere (flow rate of 50 mL/min).

## 3. Results

### 3.1. Electrospinning

Tests were conducted by varying the amount of essential fatty acids (EFA) as described in Table 1 summarizes the observations of the influence of incorporating EFA into the standard PCL polymeric solution. Environmental parameters (Temperature (°C) and Humidity (%)) were also observed, as well as the applied Voltage (kV) in the solution. These parameters are determined at the time of each test.

It was possible to prepare electrospun membranes with all the tested concentrations, with stability, and without jet dripping. The membranes containing 0.25g and 0.50g did not show beads, while those containing 0.75g and 1.00g presented visible beads on their surface, indicating oil accumulation. Additionally, it was observed that the applied voltage during the electrospinning of each membrane increased with the increase of EFA concentration in the solution. The humidity during the processes ranged from 45% to 49%. Therefore, it was defined that the maximum EFA concentration in the membrane observed in this study was 1g (representing a mass ratio of 1:1 of EFA to PCL).

**Table 1.** Study of the maximum concentration of EFA in the membrane.

According to Table 2, it was possible to prepare electrospun membranes with quantities of 0.1mL, 0.2mL, 0.3mL, and 0.4mL of SH (heparin solution). The membranes containing 0.1 mL and 0.2 mL of SH showed stability of the Taylor cone without jet dripping, and no beads were visually observed. Despite presenting stability of the Taylor cone without jet dripping, the membrane containing 0.4 mL of SH showed visible beads on its surface. This may indicate an accumulation of SH or may have occurred due to the low solubility of SH in the PCL solution.

**Table 2.** Study of the maximum concentration of SH in the membrane.

The test with 0.5mL of SH did not yield an efficient result for membrane formation due to the appearance of beads of approximately 2mm, along with the instability of the Taylor cone and jet dripping, which could potentially leave solvent residues on the membrane, making it unsuitable for further studies. This means that the maximum quantity supported by the membrane in this study was 0.4mL of SH, corresponding to 400 IU

### 3.2. Sterilization using ozone

The sterility test demonstrated the absence of colony formation or microbial growth in any of the culture media during the 7-day incubation period of the samples. Thus, the absence of contamination by microorganisms (bacteria and fungi) in the membranes after the sterilization process using O_3_ is confirmed. Consequently, it can be stated that microbiologically, ozone was effective in sterilizing the electrospun polymeric membranes in this study.

### 3.3. Morphological characterization

Figure 1 (a) shows the micrograph of PCL fibers, where a uniform and randomly distributed appearance is observed, with no beads in the fibers. The histogram (b) displays the average diameter (DM = 1.160 ± 0.9992 μm), with the maximum diameter observed being 4.849 μm and the minimum diameter being 0.198 μm.

**Figure 1.**
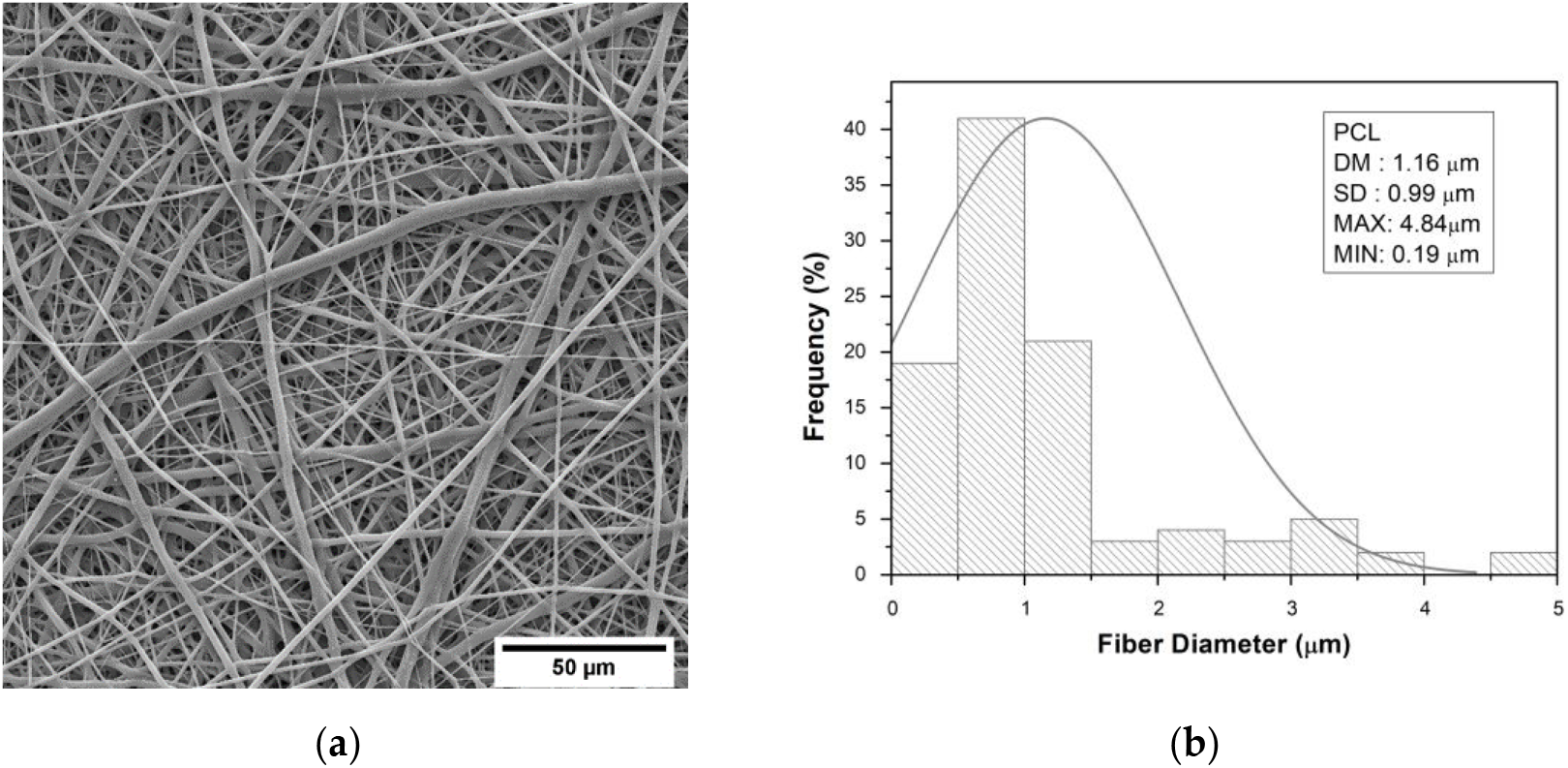
(**a**) Micrograph of PCL fibers; (**b**) The histogram presents the distribution, average diameter, standard deviation, maximum, and minimum measurements of fiber sizes, while the Gaussian curve illustrates the normal distribution of diameters around the mean value.

Figure 2 (a) displays the micrograph of PCL+SH fibers, which exhibit well-formed and randomly distributed fibers with varying diameters. However, it shows smaller diameters than those observed in the pure PCL membrane, as evident in the histogram in Figure 2 (b), with an average diameter (DM) of 0.981 ± 0.663 μm.

**Figure 2.**
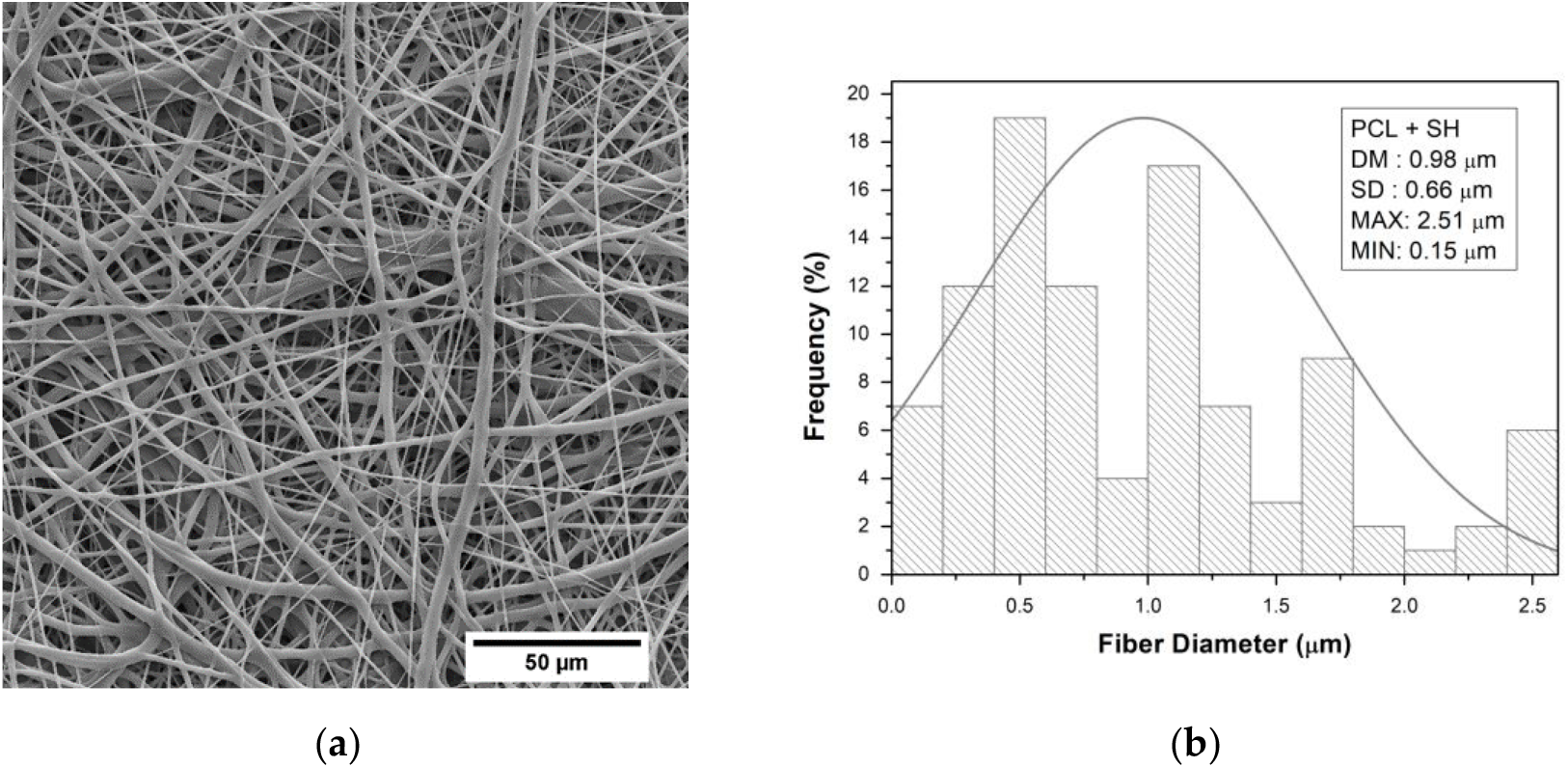
(**a**) Micrograph of PCL + SH fibers; (**b**) The histogram presents the distribution, average diameter, standard deviation, maximum, and minimum measurements of fiber sizes, while the Gaussian curve illustrates the normal distribution of diameters around the mean value.

Figure 3 (a) shows the micrograph of PCL+EFA fibers, which exhibited beads and oily residues on their surface. It can be observed that, in addition to the formation of uniform fibers, there are areas with deformations, likely caused by the concentration of oil present. In addition to the formation of uniform fibers, it can be observed that there are areas with deformations, likely caused by the concentration of oil in the membrane. According to the histogram presented in Figure 3 (b), the fibers of this material had an average diameter (DM) of 1.53 ± 0.924 μm.

**Figure 3.**
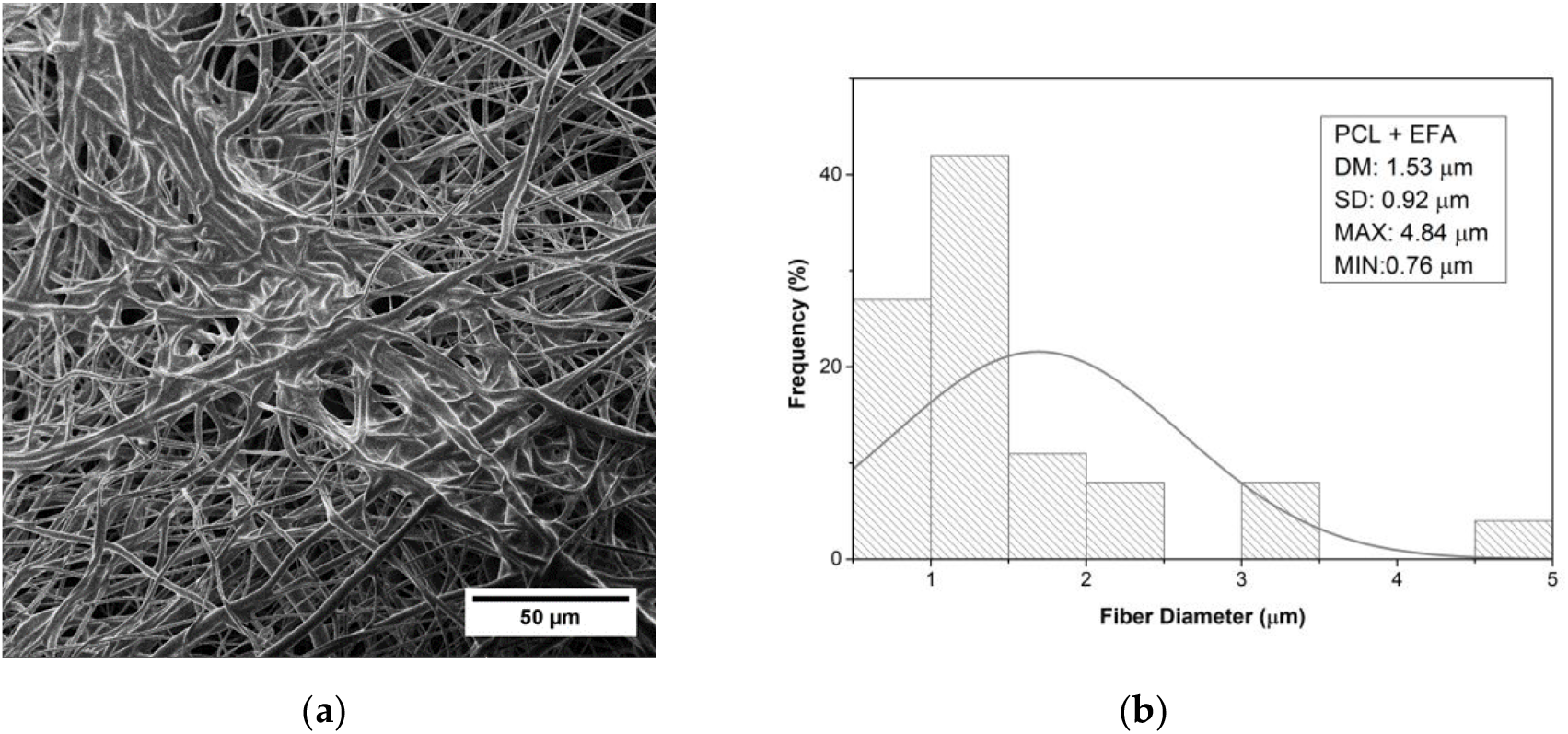
(**a**) Micrograph of PCL + EFA fibers; (**b**) The histogram presents the distribution, average diameter, standard deviation, maximum, and minimum measurements of fiber sizes, while the Gaussian curve illustrates the normal distribution of diameters around the mean value.

### 3.4. Study of membrane wettability through contact angle

Figure 4 (a) shows the contact angles of electrospun PCL, (b) PCL + SH, and (c) PCL + EFA membranes obtained for the study of wettability. The PCL membrane exhibited a contact angle of 123°, while the PCL + SH and PCL + EFA membranes showed contact angles of 102° and 95°, respectively.

**Figure 4.**
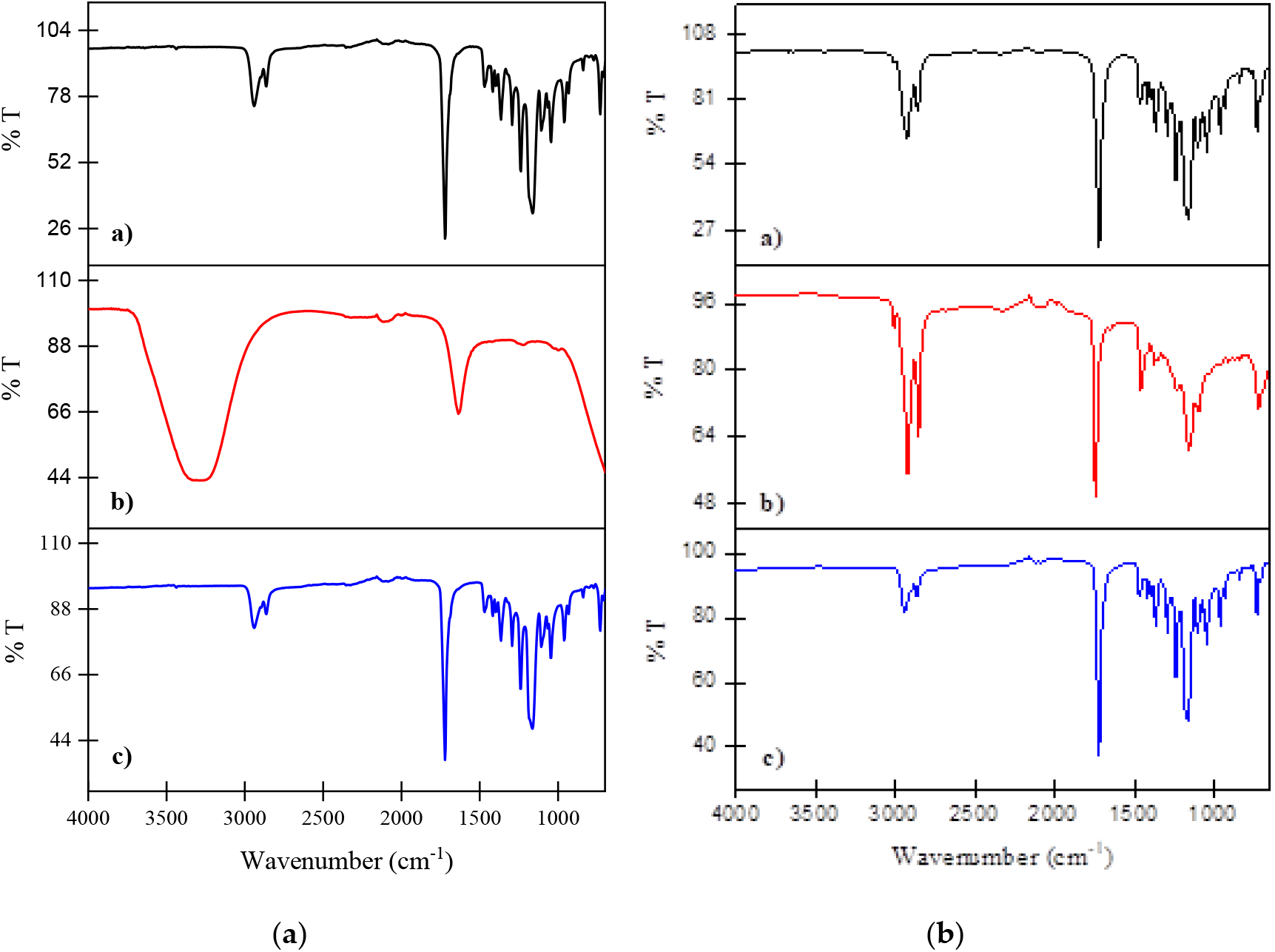
Water droplet on the (**a**) PCL membrane; (**b**) PCL+SH membrane; (**c**) PCL+EFA

### 3.5. Chemical characterization by Fourier-transform infrared spectroscopy

Figure 5 illustrates the FTIR spectra acquired from the mid-infrared region (4000 - 650 cm-1) for the following samples: PCL, Sodium Heparin, EFA, PCL with SH, and PCL with EFA.

**Figure 5.** FTIR Spectra of **(a)** a) Polycaprolactone (PCL), b) Sodium Heparin, c) Polycaprolactone (PCL) with Sodium Heparin, and **(b)** a) Polycaprolactone (PCL), b) EFA and c) Polycaprolactone (PCL) with EFA.

### 1.6. Thermal analysis of the membranes through Thermogravimetry

Figure 5 presents the curves obtained from thermogravimetric analysis (TGA) and derivative thermogravimetric analysis (DTGA) of PCL (a), PCL+SH (b), and PCL+EFA (c) membranes. No significant mass loss was observed until approximately 330°C, indicating the absence of water in the material and the absence of residues from volatile solvents used to prepare the polymer solution.

Figure 5. Water droplet on the (**a**) PCL membrane; (**b**) PCL+SH membrane; (**c**) PCL+EFA

The thermogravimetric analysis demonstrated that the incorporation of SH and EFA did not modify the thermal stability of the polymer. Still, a slight increase in the starting and ending temperatures of the thermal event was observed compared to PCL. The starting and ending temperatures of the samples are described in Table 3.

## 4. Discussion

### 4.1. Electrospinning and morphological characterization

Electrospinning is a process used to produce fine and continuous fibers from a polymeric solution. The addition of components to the solution can affect the increase in tension during the electrospinning process, which may lead to changes in the diameter of the fibers and the appearance of formed pearls.

In the context of electrospinning, adding components to the polymeric solution can affect the surface tension of the solution, which, in turn, can influence the diameter of the produced fibers. Studies on the composition of human milk, for example, show that emulsifying lipids and proteins can affect the physicochemical properties of milk fat.^13^ These properties can influence fiber formation during the electrospinning process.

Additionally, adding components to the solution can affect the appearance of pearls formed during electrospinning. Studies on the composition of human milk show that emulsified lipids in specific-sized globules can affect the stability and appearance of the formed pearls.

However, it is important to note that no specific studies were found to investigate the relationship between adding sodium heparin and essential fatty oils to the electrospinning solution and increasing tension, fiber diameter, and appearance of formed pearls. Therefore, further research is needed to better understand this relationship.

In summary, adding components to the electrospinning solution can affect the increase in tension, fiber diameter, and appearance of formed pearls. Studies on the composition of human milk show that emulsifying lipids and proteins can influence these properties. However, more research is needed to investigate the relationship between adding sodium heparin and essential fatty oils to the electrospinning solution and these parameters.

### 4.2. Sterilization using ozone

Ozone has been extensively studied for its effectiveness in membrane sterilization. Numerous studies have investigated the inactivation of bacteria and resistance genes by ozone in laboratory experiments and full-scale wastewater treatment. These studies have examined the impact of ozone on bacterial cultivability, membrane damage, and disruption of intracellular genes. It has been observed that ozone doses feasible for full-scale application can lead to the disruption of intracellular genes in bacteria, although flocs in wastewater may interfere with this effect.^14^

The mechanism of bacteria inactivation by ozone involves the destruction of cell membranes, leading to lysis reactions.^15^ Ozone treatment has been shown to cause cell membrane damage and increase membrane permeability in bacteria. The leakage of cellular components such as potassium, magnesium, and adenosine triphosphate (ATP) has been used to indicate ozone-induced damage to the cytoplasmic membrane. Ozone treatment can result in the coagulation of cytoplasm and the occurrence of agglutinations in treated cells.^16^

Ozone gas sterilization has also been investigated for its impact on electrospun scaffolds’ properties and cell compatibility. Studies have evaluated the suitability of ozone gas for sterilizing scaffolds made of polycaprolactone (PCL), a polymer commonly used in tissue engineering and regenerative medicine applications. Ozone gas sterilization has successfully sterilized the scaffolds while preserving most of their initial attributes, although some changes in mechanical properties may occur.^12^

In terms of practical applications, ozone is often used as a sterilization method due to its oxidative properties and ease of decomposition. It has been used for sterilizing medical equipment and eliminating bacteria in the soil.^17^ In wastewater treatment, ozone is typically paired with chloramines to inhibit biofilm growth on membranes, with ozone reducing flux decline and chloramines mitigating biofouling by dissolved organics.^18^

Overall, studies have demonstrated the potential of ozone for membrane sterilization. Ozone can effectively kill bacteria and other microorganisms by increasing cell membranes’ permeability and causing cell content leakage. However, the energy flux delivered to the electrodes and the ozone concentration are important factors that must be considered in sterilization. Further research is needed to optimize the use of ozone for membrane sterilization and to understand its effects on the properties of sterilized materials.

### 4.3. Study of membrane wettability through contact angle

PCL is a material characterized by low levels of surface free energy, in other words, it is hydrophobic, so it was expected that the contact angle would be greater than 90°. In the literature, authors report contact angles of electrospun PCL with randomly deposited fibers ranging from 122° to 145°. One of the causes of this variation in the contact angle of PCL is the diameter of the fibers. Additionally, a wide variation in the molecular weight of the polymers can indirectly influence these results.^19^

Regarding the PCL + HS membrane, despite the relatively low concentration of HS (400 IU), a significant modification of the surface was observed. There was a decrease in the angle of 22° compared to the PCL membrane. As mentioned, sodium heparin is water-soluble and is characterized as a hydrophilic substance, which can interfere with the modification of this material, increasing the interaction of the membrane surface with the water droplet. During this investigation, no similar materials to this membrane were found in the literature.

Regarding the angle obtained by the PCL+AGE membrane, a more contextualized discussion arises due to the specific characteristics of oily samples, such as the surface tension and polarity of the sunflower oil rich in AGE, which was used to produce this membrane. The obtained contact angle suggests that its surface has greater hydrophilicity; however, it is known that water and oil have opposing molecular properties, polar and nonpolar, respectively. This phenomenon can be seen when comparing the shape of an oil droplet in water and an oil droplet in contact with air. In contact with air, the oil molecules tend to form a spherical shape, as they will have a smaller surface area, meaning a smaller number of oil molecules in contact with the air. The oil droplet spreads in water, increasing the surface area in contact with water. A similar behavior was observed by Moraes Segundo,^19^ who produced electrospun PCL membranes with variations in the concentration of copaiba oil, where the membrane with 25% wt of copaiba oil presented a contact angle of 95°, a difference of 47° compared to the pure PCL membrane.

### 4.5. Chemical characterization by Fourier-transform infrared spectroscopy

For the FTIR spectra of PCL, it is possible to observe peaks corresponding to characteristic stretching vibrations of the material, such as in the region of 2869 cm^-1^, which corresponds to CH stretching, and a peak at 1722 cm^-1^ characteristic of carbonyl (C=O) stretching. The band in the region of 1242 cm^-1^ represents the presence of ester groups. ^20^

At 3340 cm-1, a stretching band corresponds to the O-H bond of COOH. At 2929 cm^-1^, a stretching band is related to the C-H bond. The peak at 1625 cm^-1^ is characteristic of the C=O bond. Additionally, the peak at 1417 cm^-1^ corresponds to the stretching of the C-O bond. Furthermore, it is possible to observe a peak at 1221 cm^-1^, associated with the existing C-O-C group bond, and a peak at 791 cm-^1^ corresponding to the N-H group.^20^

For the spectrum of the EFA sample, the bands in the 3011 – 3005 cm^-1^ region are attributed to oleic and linoleic acids. Values at 3006 and 2851 cm^-1^ were observed, corresponding to the symmetric and asymmetric C-H vibrations in CH2 and CH3 groups. The bands around 1745 cm^-1^ are assigned to the ester carbonyl.^21^

The bands at approximately 1654 cm^-1^ are related to the C=C bonds in unsaturated fatty acids. Bands between 1445 – 1440 cm^-1^ are characteristic of C-H vibration or deformation. The peak at 1376 cm^-1^ corresponds to the deformation vibration of the methylene group, and the one at 1233 cm^-1^ corresponds to the deformation vibration in the plane of the =CH group, which represents non-conjugated double bonds.^21^

Bands in the 1300 – 1000 cm^-1^ range are characteristic of the C-O ester linkage. The bands corresponding to C-C bonds can be found in 1100 – 1000 cm^-1^ and 900 - 800 cm^-1^. The C=O bond has characteristic bands at 1159 cm^-1^ and 965 cm^-1^. The absorption band around 721 cm^-1^ is attributed to the (CH2) group. When analyzing the spectrum of the PCL membrane with EFA, it is possible to observe bands related to oil contributions, which are: 3006 cm^-1^ (-OH), 2851 cm^-1^, 1745 cm^-1^, 1654 cm^-1^ (C=C), 1378 cm^-1^, 1233 cm^-1^ (=CH), and 1159 cm^-1^ (C=O).^21^

With the analysis of the infrared spectrum of sodium heparin, the presence of peaks corresponding to the functional groups in its chain is observed, which, according to Coelho, are the main peaks for HS, such as the OH, CH, NH, and C=O groups. The main bands are the 3340 cm^-1^ stretching band of the COOH O-H bond, and 2929 cm^-1^ referring to the C-H stretching band; 1625 cm^-1^ characteristic band of the C=O bond, and 1417 cm^-1^ axial deformation of the C-O group; 1221 cm^-1^ asymmetric axial deformation of the C-O-C group; 791 cm-1 for the out-of-plane N-H group.

However, when analyzing the spectrum of PCL+HS membrane, the small contribution of HS is evident only in the 1417 cm^-1^ band, as the bands at 2929 cm^-1^, 1221 cm^-1^, 1028 cm^-1^, and 791 cm^-1^ coincide with the bands shown in the PCL spectrum.^22^

### 4.6 Thermal analysis of the membranes through Thermogravimetry

According to Mohammed^23^, the onset temperature of PCL degradation is around 340°C, and the final temperature is 489°C. Regarding SH, Coelho^22^ presents the onset temperature of SH at 260°C and the final temperature at approximately 600°C, and finally, Rodrigues and Weber^24^ show that the onset temperature of EFA is about 300°C, with a maximum temperature of 500°C. Thus, the temperatures reported in the literature are close to the values obtained in this study.

The mass losses are observed at: 430.18°C with a mass loss of 49.98% for the PCL membrane, 435.65°C with a mass loss of 50.04% for the PCL+SH membrane, and 437.03°C with a mass loss of 50.02% for the PCL+EFA membrane. As exposed, the interaction between the drugs and PCL did not significantly modify the material’s thermal stability.

## 5. Conclusions

The membranes of Pure PCL, PCL+SH, and PCL+EFA were successfully obtained by electrospinning.

Using ozone gas for sterilization proved effective since the sterility test showed no contamination by microorganisms (bacteria and fungi), making the membranes suitable for biomedical applications. The morphology of the membranes revealed that the obtained fibers were uniformly and randomly distributed. The statistical study of fiber diameter showed that adding drugs altered the fiber diameter. The average diameter of Pure PCL fibers was DM = 1.160 ± 0.992 μm, while the average diameter of PCL+SH fibers was DM = 0.981 ± 0.663 μm, and PCL+EFA fibers had a diameter of DM = 1.53 ± 0.924 μm. Regarding the wettability of the membranes, it can be stated that both SH and EFA modified the membrane surface, altering the contact angle. The contact angles for PCL, PCL+SH, and PCL+EFA membranes were 123°, 102°, and 95°, respectively. FTIR analysis of the PCL+SH membrane spectrum showed a small contribution of SH in the 1417 cm^-1^ band, corresponding to the C-O bond. In the spectrum of the PCL+EFA membrane, bands related to the contributions of the oil were observed, which were: 3006 cm^-1^ (-OH), 2851 cm^-1^, 1745 cm^-1^, 1654 cm^-1^ (C=C), 1378 cm^-1^, 1233 cm^-1^ (=CH), and 1159 cm^-1^ (C=O).

Thermogravimetric analysis of the membranes revealed no substantial modification in the thermal stability of the membranes due to the interaction between SH and EFA with PCL. The membranes developed in this study show high potential for application as dressings and drug delivery systems. Incorporating other substances can impart even more attractive characteristics to these materials. Obtaining blends with other polymers, such as cellulose acetate, chitosan, and hyaluronic acid, among others, may modify the thermal and mechanical properties of the membranes. Additionally, including metallic nanoparticles, such as gold and silver, is of particular interest due to their important properties for medical applications, as these materials exhibit antibacterial and anti-inflammatory activity, providing significant benefits in wound treatment.

## Author Contributions

Conceptualization, W.B.; T.F. and J.R.; methodology, T.F.; formal analysis, K.S.; investigation, T.S.; J.C and I.B.; resources, Y.B.; data curation, C.R.; writing—original draft preparation, T.F.; writing—review and editing, J.S.J.; supervision, M.P. and W.B.; All authors have read and agreed to the published version of the manuscript.

## Funding

This research received no external funding

## Data Availability Statement

Data will be available upon request.

## Acknowledgments

Our acknowledgments to Universidade Federal do Amazonas (UFAM), Fundação de Amparo à Pesquisa do Estado do Amazonas (FAPEAM) and Coordenação de Aperfeiçoamento de Pessoal de Nível Superior (CAPES).

## Conflicts of Interest

The authors declare no conflict of interest.

## Notes

### Competing Interest Statement

The authors have declared no competing interest.

